# Native language leaves distinctive traces in brain connections

**DOI:** 10.1101/2022.07.30.501987

**Authors:** T. Goucha, A. Anwander, H. Adamson, A. D. Friederici

## Abstract

The world’s languages differ substantially in their sounds, grammatical rules, and expression of semantic relations. While starting from a shared neural substrate, the developing brain must therefore have the plasticity to accommodate to the specific processing needs of each language. However, there is little research on how language-specific differences impacts brain function and structure. Here, we show that speaking typologically different languages leaves unique traces in the brain’s white matter connections of monolingual speakers of English (fixed word order language), German (with grammatical marking), and Chinese (tonal language). Using machine learning, we classified with high accuracy the mother tongue based on the participants’ patterns of structural connectivity obtained with probabilistic tractography. More importantly, connectivity differences between groups could be traced back to relevant processing characteristics of each native tongue. Our results show that the life-long use of a certain language leaves distinct traces in a speaker’s neural network.

## Introduction

All humans share the neurobiological equipment that allows them to learn the language they are born into^1,2^. Considering the universality of the cognitive infrastructure underlying language^2^, one could easily deduce that all languages should be rather similar. Yet, this stands in stark contrast with the actual scope of variation that can be observed in the languages across the globe^3^. How the human cognitive system and its neurobiological basis are able to deal with this linguistic variety still is an open question. We will approach this question by first considering the language differences and their different processing demands and then explore to what extend these differences lead to modulations in the human neural system underlying language.

The languages of the world are grouped into families according to their genealogy, that is, which ancestors they have and how long ago they diverged. For example, Italian or French are classified as Romance languages because both evolved from Latin. Together with Germanic (e.g. German and English) and Slavic languages, they belong, in turn, to the higher-order family of the Indo-European languages. Yet, some of the closely related languages within each of these families are still typologically very diverse and underwent changes both with respect to their lexicon and grammar^4,5^.

The diversity across the languages of the world expresses itself in three main language domains: phonology, concerning its externalisation in sounds (for spoken languages) or signs (for sign languages), semantics, which deals with content, and syntax, regarding how words are structured into sentences. Starting with the sound systems that underlie each language, there are virtually no limits to their possible phonetic repertoires^6^. Among spoken languages, there is a fundamental distinction between those that only use vowels and consonants to differentiate between words as in the Indo-European languages, and tonal languages, such as Chinese or Vietnamese, which also use different pitches or melodies on each syllable to distinguish words that otherwise would sound identical. Concerning semantics, the lexicon is organised according to general principles^7^, which remain considerably stable along language evolution^4,5^. Here, differences are more specific to particular topics^8,9^, especially as to how the words of a certain language reflect its particular sociocultural context^10,11^. Regarding syntax, human languages seem to follow a basic computational principle that combines words into hierarchical structures, building phrases and sentences^2^. However, the way this hierarchical structure is externalised into a sequence strongly depends on the specific language. A sentence usually describes who is doing what to whom, by saying what the subject (S) of the main verb (V) is doing to a person or object (O). Languages are classified typologically according to the preferred order in which these elements appear, the so-called canonical word order^12^, with a strong preference worldwide for either SVO (e.g., English and German) or SOV (e.g., Japanese). Additionally, languages use different cues to distinguish the subject from the object. English, for example, has a fixed SVO canonical word order, which clearly determines that the first noun phrase is the subject and the second the object. Other SOV languages like German, mark the subject and the object grammatically (e.g., by a particular word ending to convey case marking information), which allows sentence elements to move more freely in the sequence^12^. In sum, the diversity among languages and the way they convey information lead to the conclusion that the cognitive apparatus allowing us to acquire language is originally universal and open for each language, but then progressively adapts to the particular characteristics of the speaker’s mother tongue along lifetime.

In fact, psycholinguistic research focussing on language processing has shown that the complex task of acquiring language starts in the mother’s womb with learning the melody and rhythm of our native language^13^. The human new-born proceeds with fine-tuning to its particular sound repertoire^14^, which is achieved mostly within a year at the cost of a substantial loss of the ability to discriminate and learn new sounds from then on^1^. Subsequently, after having acquired their first words, children begin to combine words into bigger chunks, and eventually start building sentences. They have to recognise the different cues in their language in order to identify who did what to whom in a sentence^15^. Thus, adult speakers of different languages prioritise different information types (e.g. word order, case marking) during language comprehension^16^. Such cross-linguistic processing differences can be observed in the brain activity of speakers of different languages while listening to sentences with similar properties^17^. Brain imaging studies often stressing commonalities across languages^18,19^, also report cross-linguistic differences in brain activity^20^. In conclusion, not only do languages vary strongly in the way they are organised, speakers seem to adapt to such characteristics when processing their mother tongue.

If so, one would expect that such differences would be traceable in the human brain. The brain is known to generally adapt to its environment during development^21^, and connections in the brain can undergo extensive rewiring even in the adult brain^22^. This can be achieved by strengthening the connections in stronger use^23^, which makes the conduction of the neuronal signal more efficient, while losing those that become obsolete. Most studies investigating brain plasticity so far focus on the short-term effects of an experimental intervention involving a specific task^24^, a lifelong scale such as the use of a particular language processing strategies should reveal observable effects in the brain. This hypothesis is further motivated by the fact that the fibre pathways connecting the brain areas within the language network still undergo strong maturation after birth well into adolescence^25,26^, and that their maturation stage is linked to language performance^27^. It is, thus, plausible that the trajectory of the growth of these white matter pathways is influenced by the use of one’s native language.

To test this hypothesis we selected three typologically different languages that represent some of the strongest linguistic differences we have discussed above. Here, we selected English and German, two Indo-European languages of the Germanic branch that, despite being closely related, differ fundamentally in their syntactic structure. As a third and final language, we opted for Chinese, a Sino-Tibetan language which exhibits lexical tone, among other singularities introduced subsequently. In particular, we used diffusion MRI data to compare the brain structural connectivity of speakers of different mother tongues. We expected the differences in processing across languages to be structurally reflected in the white matter connections that support language processing.

Neurally, language is processed in a brain network mainly comprising brain regions around the Sylvian fissure in the left hemisphere, which are connected by fibre pathways that run either dorsally or ventrally to this anatomical landmark^28–32^, and can be partly found mirrored in the right hemisphere^33^. The posterior temporal cortex is a region where these white matter pathways overlap, being a point of convergence of dorsal and ventral processing streams^28,34^, frequently being implicated in integration of different types of information and sentence-level processing^28,34,35^. That is why we reconstructed the white matter connections that cross this region, hence obtaining a map of structural connectivity^36^ of the language regions in each participant. Additionally, this also allowed us to analyse the transcallosal connections of the temporal cortex connecting the two hemispheres, which were shown to be crucial for the processing of intonation and the integration of prosody in sentence comprehension^37,38^. As a novel approach in language studies, we used machine learning^39^ to assess whether the overall pattern of connectivity of each participant contained information to infer their respective native language. After establishing that the connectivity maps allowed an accurate classification, we set out to identify the regions of the language network showing a significant modulation of their connectivity according to one’s mother tongue. By comparing the connectivity maps of the three groups, we could assess whether the processing differences between the three languages corresponded to differences in structural connectivity. Concerning our hypothesis, we addressed the differences between each of the languages. We started to consider the differences between English and German. German – as already mentioned – is a language with free word order and is highly marked by grammatical cues^40–42^, which are used by speakers to retrieve the sentence structure^15–17,43^. These processes were shown to recruit the left inferior frontal gyrus (IFG)^28,34,40^, at the frontal end of the dorsal pathway^27,28^, which deals with the abstract sentence structure that is inferred from grammatical rules^44^. English sentences mostly have a reliable, fixed word order^15,41^, and speakers of English are more influenced by semantic cues, such as animacy^16^, or meaning associations between sentence elements, which are mainly processed in the ventral stream^29,30^. For this reason, we expected stronger dorsal connectivity to the inferior frontal cortex in the German-speaking group, whereas the English-speaking should in turn display stronger connectivity in the ventral stream. Concerning Chinese, we first focus on the differences between tonal and non-tonal languages. To date a number of studies have shown a more bilateral involvement while processing lexical tone in both spoken^45^ and written^46^ language, in comparison to atonal languages^20,47^, which was also shown to be reflected in brain morphology^48^. Although the lexico-semantic processing of lexical tone is mainly left-lateralised, the processing of pitch information is in general less lateralised than the processing of the acoustic features that distinguish other speech sounds as consonants and vowels^49^. Additionally, the pitch information from lexical tone must be integrated with sentence prosody, known to be processed in the right hemisphere^33^. Altogether, this would require a stronger cross-talk between both hemispheres in Chinese speakers. That is why we expected stronger connectivity in the right hemisphere and in the fibres of the corpus callosum in this language group. Besides, Chinese is known for its exceptionally large number of homophones^20^, even when taking lexical tone into account. These three languages also differ in other aspects, for example in their writing systems and orthography depth, which will be discussed in more detail later^19^. In sum, we hypothesised that the specific processing demands from each of these three typologically different languages would lead to differences in the strength of the white matter fibre pathways of their speakers. Here, we show that this is indeed the case by using a multivariate pattern recognition analysis on the structural connectivity of two independent samples of speakers for each of the three languages, followed by a mass univariate analysis to localise those differences.

## Results

### Mapping connectivity of language regions with fibre tractography followed by classification with machine learning

Using probabilistic tractography^36^ with seeds in the posterior superior and middle temporal gyri (pSTG, and pMTG respectively), we mapped the structural connectivity in a total of 134 monolingual native speakers of Chinese, English, and German, with two independent subsamples for each language, matched for sex, age, and education (Chinese: N = 30 + 18; English: 20 + 18; German: 30 + 18). First, we were able to consistently map a universal network of white matter pathways connecting the brain regions typically involved in language processing, which was common to all subjects in both subsamples of each language (Supplementary Figure S1). The brain images of all subjects were registered to a balanced sample-specific template to minimise potential effects of any population differences in global brain morphology. First, we did not find significant differences in brain volume across groups. Additionally, to prevent further analyses from yielding results in regions with systematic differences in brain shape between groups, we excluded all areas requiring strong deformation during normalisation to our template (Supplementary Figure S2). We then applied machine learning to classify the connectivity maps we obtained for each language data set. We trained a Gaussian process classifier in a k-fold cross-validation scheme on the whole dataset, which was able to predict the language corresponding to the brain structure in the test data-set with high accuracy. The classifier performed significantly above chance (p < 0.001, assessed by a permutation test with 10,000 permutations) with classification accuracies ranging from 68.64% to 76.46%. Figure 1 graphically displays the performance of the Gaussian process classifier in the left hemisphere (Supplementary Figure S3 corresponds to the analogous analysis performed in the right hemisphere). Additionally, to assess whether this result was generalizable between independent datasets with different scanning conditions, we conducted a prediction analysis by training the classifier in the first and bigger subsample of matched participants (N = 80) to predict the language of the second subsample on the basis of their connectivity (N = 54). The classifier still performed significantly above-chance accuracy for three of the seed ROIs (left pSTG, right STG, and right MTG), with accuracies ranging 55.12-61.11% (p < 0.01, assessed with permutation test). We finally ran the classifier only within the datasets acquired in the same scanner at the same site (48 German speakers, and 18 Chinese speakers), to exclude a scanner-specific effect, and its performance remained above chance (p < 0.01, assessed with permutation test, taking into account uneven sample sizes).

**Figure 1.**
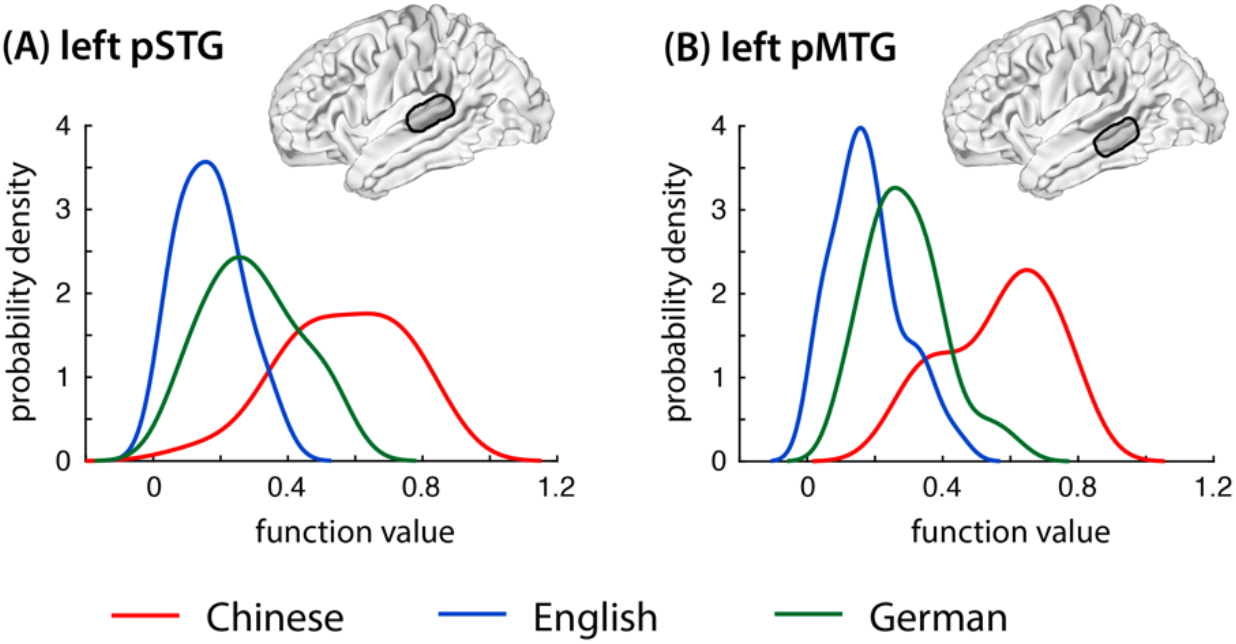
Performance of the classifier. Performance of the classifier on the connectivity map of the two temporal seed ROIs in the left hemisphere, (A) the left posterior superior temporal gyrus (pSTG) and (B) the left posterior middle temporal gyrus (pMTG). Performance for the three different languages is colour coded red for Chinese, blue for English and green for German.

### Localisation of connectivity differences using voxel-wise analysis

After showing that the connectivity maps contained information to decode the mother tongue of a subject accurately, we proceeded to localise the regions with significant differences in structural connectivity across the three groups, using a conventional mass univariate approach. More concretely, we performed spatial voxel-wise statistical comparisons of the connectivity maps of the speakers of each language. As one would expect from the previous results, the white matter network, while fundamentally shared between all participants, showed locally specific modulations by each of the three languages. Once again, areas requiring strong deformation in the registration to the template space were excluded.

### Connectivity differences from seed ROIs in the left hemisphere: Conjunction analysis

To summarise the main findings concerning regional differences in brain connectivity across the three languages, we first present the result of a conjunction analysis for the sake of conciseness. Figure 2 therefore shows brain areas where one language group displayed significantly higher connectivity values than either of the other two languages across the two subsamples (see Supplementary Figure S4 for slice views). However, the results from the direct pair-wise comparison of the connectivity maps of each of the three pairs of languages are largely superposable (and are thus exhaustively presented in the next section). In particular, German speakers exhibited a cluster with stronger connectivity between the pSTG with the IFG via the dorsally located arcuate fascicle. English speakers, in turn, showed a stronger connectivity of the pMTG with a cluster in the anterior temporal cortex via the ventrally located inferior and middle longitudinal fascicles and extreme capsule fibre system. Finally, Chinese speakers displayed stronger connectivity of the left pSTG to the right hemisphere, with clusters spanning through the corpus callosum to the contralateral temporal cortex and another cluster reaching into the subcortical grey matter, especially at the thalamus. Furthermore, the left pMTG in this group similarly presented stronger connectivity to cortical and subcortical regions in the right hemisphere via the corpus callosum. Additionally, we found connections in the left hemisphere to the contiguous *planum temporale* extending to the parietal cortex and ipsilateral subcortical grey matter regions. In sum, the group comparison of the connectivity maps across the three languages demonstrated significant differences along the white matter pathways of different processing streams (Figure 2, Supplementary Figure S5, Supplementary Figure S6).

**Figure 2.**
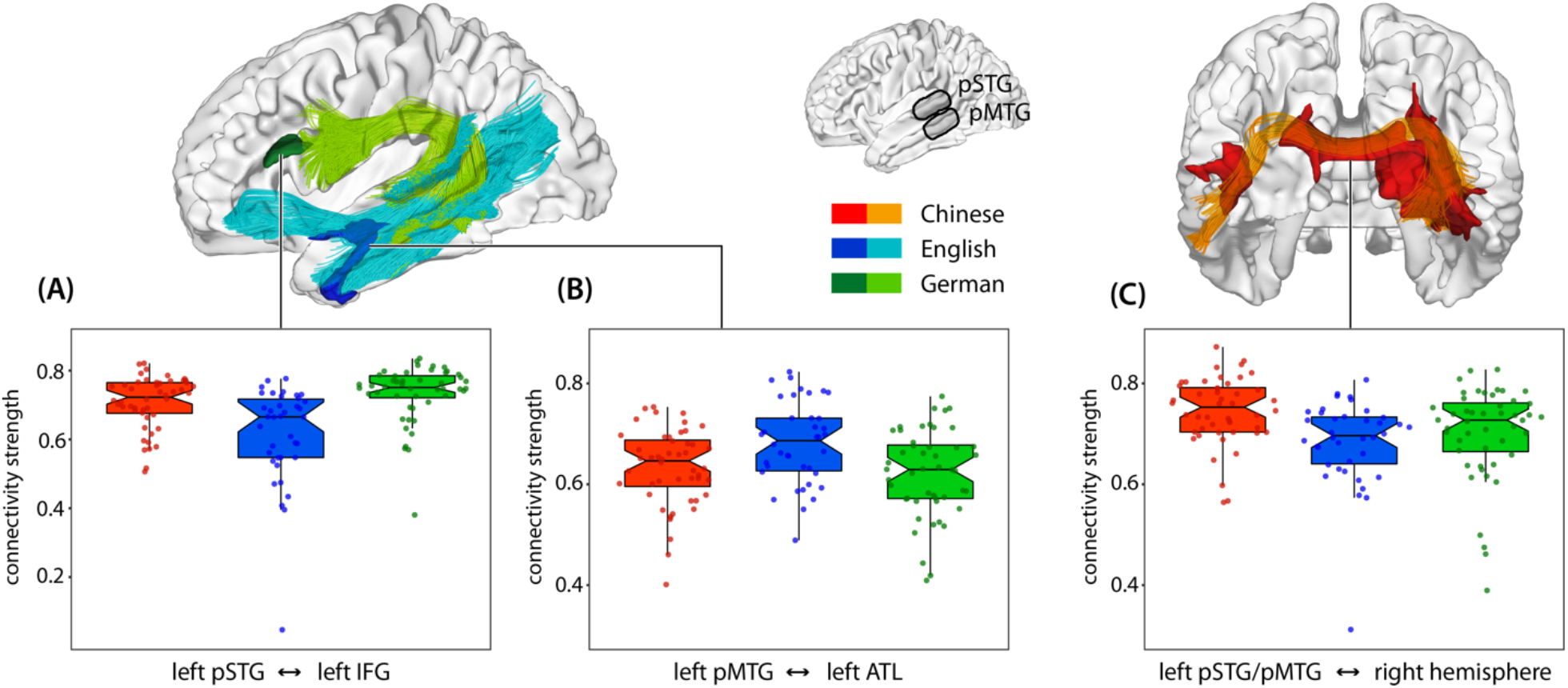
Cross-linguistic differences in white matter connectivity. White matter connectivity from left Wernicke’s area (posterior superior/middle temporal gyrus, pSTG, pMTG) to (A) the left inferior frontal gyrus (IFG), and (B) the left anterior temporal lobe (ATL) and to (C) the right hemisphere. Conjunction analysis on the seed regions in the left hemisphere. The connectivity strength for the three languages is colour coded in red for Chinese, blue for English and green for German. The box-plot shows median, quartiles, 1.5x interquartile range and all individual data points.

### Connectivity differences from seed ROIs in the left hemisphere: Pair-wise comparison

A more detailed pair-wise direct comparison between each two languages corroborated the previous results (Figure 3). First, the comparison between English and German speakers yielded higher ventral connectivity from the pMTG to the anterior temporal lobe in the English group, as opposed to higher dorsal connectivity along the arcuate fascicle from both the pSTG and the pMTG to the inferior frontal gyrus in the German group. Furthermore, the pSTG seed in the German group displayed higher bilateral subcortical connectivity and transcallosal connectivity in premotor regions. Second, when comparing Chinese and English speakers, the former showed stronger inter-hemispheric connectivity with the contralateral temporal lobe, at the corpus callosum in the frontal and temporal regions from both the pSTG and pMTG. Additionally, both the pSTG and pMTG seeds displayed higher connectivity along the dorsal stream to the IFG. The English group showed, in turn, no significant clusters with stronger connectivity. Finally, when comparing Chinese and German speakers, Chinese speakers exhibited a stronger connectivity between both the pSTG and the pMTG and the cortical and subcortical contralateral regions. Additionally, the pMTG in Chinese speakers showed stronger connectivity to a cluster in the left pSTG. In turn, the German group showed no significant clusters with stronger connectivity from the left pSTG or pMTG. In the right hemisphere, we mainly found stronger transcallosal connectivity to the left hemisphere in the Chinese group, in line with our findings for this language in the left hemisphere (see Supplementary Figure S7 and Supplementary Figure S8 for more detail).

**Figure 3.**
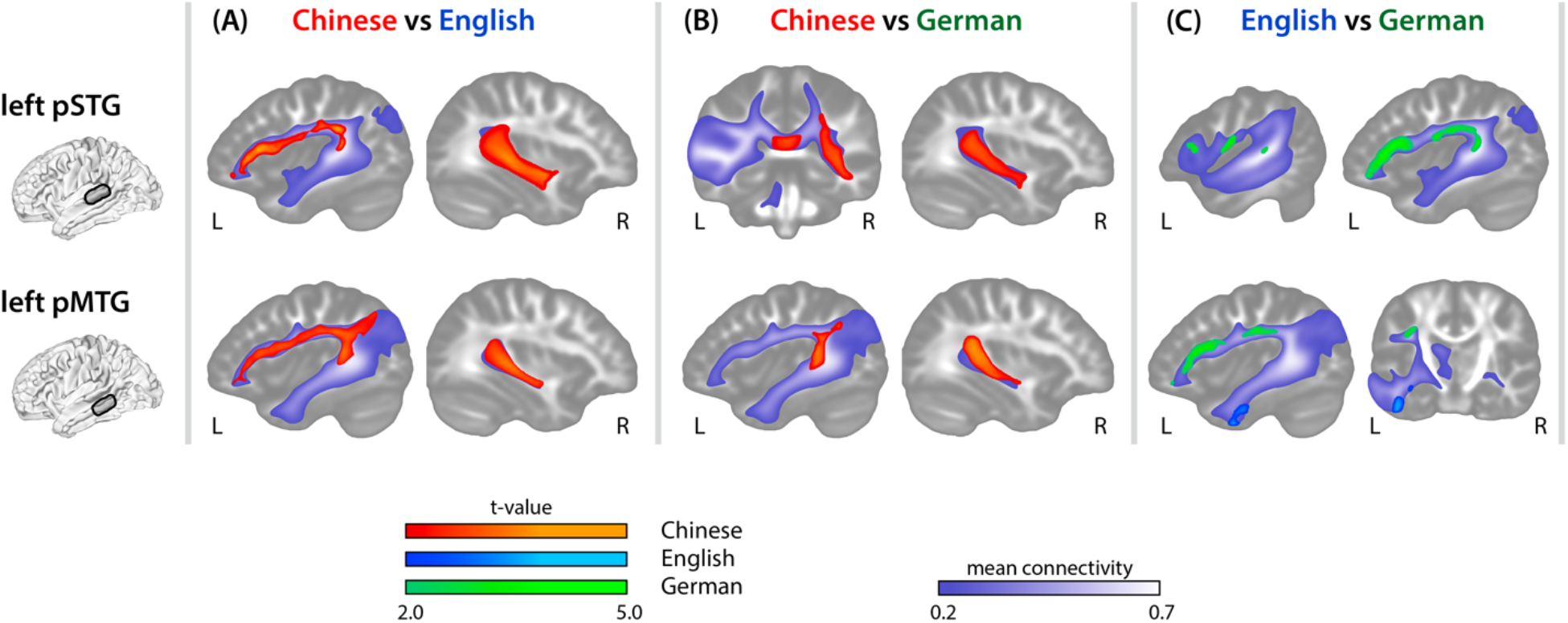
Pair-wise differences in connectivity. Connectivity differences between each pair of the three languages from left seed regions (pSTG and pMTG), (A) Chinese versus English, (B) Chinese versus German, (C) English versus German.

## Discussion

Our study provides evidence that the life-long use of a particular language leaves a distinct footprint in the brain’s structural connectivity. We showed that the individual connectivity pattern was sufficient to accurately classify the specific mother tongue of each participant in our sample. We found that although the major white matter fibre pathways comprising the language network are present in all participants, the strength of the connections along the different neural pathways is modulated according to the specific characteristics of the speaker’s mother tongue. Specifically, the results indicate that different processing demands of a given language leave particular traces in the white matter language network. Cross-linguistic differences have been reported earlier in behavioural studies^3,9,50^ and electrophysiological studies^17^ for phonological and lexical processing, and moreover, for how grammatical rules of a language relate to the way a language conveys information^17,51,52^. Even though there are universal principles guiding language acquisition^53–56^, the languages of the world provide their users with different cues to retrieve the underlying structure of a given sentence^15,17^. Here, we demonstrated that three languages which have different cues and correspondingly imply different processing demands lead to differences in the brains’ structural connectivity within the language network.

The connectivity differences we found at the brain level are congruent with the specific processing demands proposed for each language investigated in this study. First, we demonstrated a clear contrast between the dorsal and ventral pathways between English and German speakers. Although both languages belong to the Indo-European family, English has fairly scarce grammatical marking, while German uses grammatical markers to convey the relations between sentence elements^41^. Accordingly, German speakers resort to the grammatical cues during sentence processing^15,17^, implying higher demands in application of syntactic rules concerning sentence structure. Here we showed that such demands give rise to a stronger recruitment of the arcuate fascicle in the dorsal pathway – a white matter fibre pathway that has been correlated with processing complex sentence structures^27,44^. English speakers, in contrast, resort less to the infrequent cues concerning grammatical marking^15,44^, comparatively depending more on semantic information to infer the content of an utterance. We showed that at the neural level, English speakers more strongly engage the ventral pathway – a pathway which has been associated with language comprehension and in particular semantic processing^30,57,58^. The comparison between German and English speakers shows that two languages that belong to the same language family, but differ in their processing demands, influence the white matter brain structure differentially.

Chinese, in contrast to English and German, is a tonal language belonging to the Sino-Tibetan family. This implies that Chinese requires steady tracking of pitch information, partly processed in the right hemisphere, and phonological and lexical information processed in the left hemisphere. The stronger white matter connection between the two hemispheres via the posterior part of the corpus callosum we found in Chinese speakers compared to German speakers is taken to reflect the stronger bilateral involvement shown for processing tonal languages^19,47^ and the transcallosal connectivity which is the basis for the integration of pitch information with other linguistic information^37,38^.

German, English and Chinese also differ from each other in their writing systems. Whereas Chinese is logographic both English and German are alphabetic languages with German having a very shallow orthography with an almost direct correspondence between graphemes and phonemes, while English has a rather deep orthography with a more opaque correspondence between how a word is written and its pronunciation. These two types of processing of alphabetic writing have respectively been associated with either a stronger engagement of the dorsal or ventral pathway^19^. In the case of Chinese there is no clear evidence as to which brain regions are preferentially recruited in reading^19,46^.

A final consideration about the cross-linguistic differences in this study regards word order and length of the dependencies established between elements in a sentence, for example, the dependency between the verb and its object. Some languages (such as German and Chinese^59,60^) have on average a much higher dependency length than other languages (such as English and most Romance languages^59^). The processing of long distance dependencies, necessary in Chinese and German, should recruit the dorsal stream given their role in sentence structure building^28^ and its connection to the inferior frontal gyrus, recruited in time-dependent reordering of sentence elements^31,61^. This hypothesis is once again in agreement with our findings concerning connectivity differences across languages and especially the stronger dorsal connectivity of German and Chinese speakers in direct comparison to English speakers. In conclusion, our results point to a link between the specific processing demands of each language and the observed differences in brain structure.

Several precautions were taken to prevent unwanted misinterpretations due to possible confounds. First, we obtained two independent samples of speakers for each of the three languages to improve generalisability of the results and ensure the differences were not sample-specific. In fact, we could train the classifier with a first subsample for each language to then accurately predict the corresponding language on the basis of the connectivity in the other subsample. Second, in each of the two subsamples the participants in the three groups were matched for socio-demographic variables, in particular age, sex, and educational background. Third, to avoid that our results could be attributed to systematic differences in brain geometry, we excluded areas with strong differences in brain shape between groups from the voxel-wise analysis. Finally, the brain regions used as seeds for tractography as well as the regions exhibiting significant group differences we found here occur in white matter fibre pathways with a major role in language processing^27,30,62^ and their differences can be explained by the specific processing demands of the languages under investigation.

Moreover, the present results show no anatomical overlap with imaging studies assessing social and cultural differences between Western and Eastern populations. The effects in these studies consistently involved another set of brain regions, frequently including the medial prefrontal cortex^63^, but not areas belonging to the language network. In sum, the enumerated arguments strongly indicate that the present white matter differences are indeed due to the life-long use of the respective language. Recent genetic and neurobiological data support the view that the differences in brain structure we observed result from experience rather than from strong biological predisposition^64^. If our results were a mere consequence of innate genetic differences, we would expect that geographical proximity^65^ should strongly determine the extent of the differences in brain structure, with the connectivity of German speakers and English speakers being very similar, and Chinese speakers with much stronger differences. However, the degree of dissimilarity in white matter connectivity in several regions between the Chinese group and both the English and the German group was comparable to the difference between the two European groups. This suggests that other mechanisms play a major role here.

Our data rather suggest that brain plasticity occurs due to the differential recruitment of parts of the brain network, putatively as an instance of activity-dependent white matter plasticity^23,66^. In fact, the white matter pathways that compose the language network are already present, but not fully developed at birth^67^, which allows them to be shaped as a function of environmental requirements. This view is additionally supported by a functional brain study^47^. The present results illustrate how plasticity in white matter^21^ allows the brain to adapt to its environment, even with respect to a higher cognitive function shared by all humans. The innate neural system with universal principles^1,55,56,68^ adapts progressively to its input^13,68^ and is ultimately shaped by it. A common genetic endowment providing the neurobiological foundations of cognition, eventually gives rise to different structures in accordance to environmental exposure. Here, we provide evidence that the systematic yet subtle life-long processing differences required by a cognitive function, namely language processing, can give rise to structural brain differences. In conclusion, the outstanding human capacity to proficiently learn the complex system of symbols and rules that constitutes a human language seems to not only lie in a neurobiological predetermined faculty, but also requires the ability of our brain to adapt to the specific demands of each language in human development.

## Author Contributions

T.G., A.A., and A.D.F. designed the research; T.G., H.A. and A.A. analysed the data; and T.G., A.A., and A.D.F. wrote the paper.

## Data and materials availability

All data needed to evaluate the conclusions in the paper are present in the paper and/or the Supplementary Information.

## Methods

### Participants

Three groups of young German, English, and Chinese speaking students were scanned in a Siemens 3T TimTrio magnetic resonance imaging (MRI) device (Siemens Healthineers, Erlangen, Germany). From these groups, we selected datasets that meet strict quality criteria of no measurement artifacts or neurological anomalies. We further matched the demographic variables gender (71 female, 63 male), age (24.8 +/−3 years), handedness^1^, and level of education between groups, resulting in a total of 134 monolingual native speakers participants of each of the three languages.

The data was acquired in two different samples covering all three languages in each study. The first sample included 30 German, 20 English and 30 Chinese datasets scanned on MRI machines of the same model in Leipzig (Germany), Cambridge (UK) and Beijing (China), with a well-matched scanning protocol^2,3^. The second sample included 18 German, 18 British and 18 Chinese datasets. The German and Chinese datasets were scanned on the same MRI device in Leipzig (Germany), and the English speaking participants were scanned on an MRI machine of the same model in Glasgow (UK) with the same protocol as the German group. The detailed scanning parameters are described in the following section. Written informed consent was obtained from all participants, and data acquisition was approved by the respective local ethics committees, the Institutional Review Board of Beijing Normal University Imaging Center for Brain Research, the ethics committee of the College of Science and Engineering at the University of Glasgow, the Cambridge Psychology Research Ethics Committee, and the Ethics Committee of the University of Leipzig.

### Imaging data acquisition, preprocessing, and generation of group-specific template

All participants obtained a T1-weighted structural image with 1 mm isotropic resolution using a 3D MPRAGE sequence with whole-brain coverage. In the first sample, in the English and German groups, diffusion MRI images where acquired using 64 diffusion directions and a b-value of 1000 s/mm^2^ and one non-diffusion-weighted image^2^ (parameters: repetition time, TR = 6.5 s, echo time, TE = 93 ms, GRAPPA acceleration factor 2, 12-channel acquisition head coil, 48 axial slices, 2.5 mm thickness, in-plane resolution=1.8×1.8 mm^2^). The diffusion MRI datasets of the Chinese participants were acquired with a very similar protocol using 64 diffusion directions with b=1000 s/mm^2^ and one non-diffusion-weighted image^3^ (parameters: TR = 7.2 s, TE = 104 ms, 49 axial slices, 2.5 mm thickness, in plane resolution= 2.0×2.0 mm^2^). In the second sample, for all three groups diffusion MRI images where acquired using 60 diffusion direction with a b-value of 1000 s/mm^2^ and seven non-diffusion-weighted images with an isotropic voxel size of 1.7 mm and a 32-channel acquisition head coil.

Preprocessing was done in a consistent way for the datasets of all three groups in both samples. The brain was segmented from the T1 weighted image and aligned with the AC-PC coordinate system^4^. The diffusion images were denoised^5^, corrected for motion and registered^6^ to the structural image using the FSL software (University of Oxford, UK). The diffusion tensor and the fractional anisotropy (FA) images were computed in the native diffusion resolution. Finally, the distribution of up to two crossing fibre directions per voxel were computed using FSL for probabilistic crossing fibre tractography^7^.

Additionally, a high-resolution FA image was created at 1 mm isotropic resolution for each participant, and a balanced sample-specific template was generated from those images using the ANTs software^8^ using a random selection of 60 participants (20 from each language group). This template was used for the definition of regions of interest (ROIs) for tractography and for the normalisation of the tractography results of each participant. This avoided having a bias towards the specific brain structure of any of the three groups. The two datasets from each sample were combined for each language group to perform statistical analysis across groups to increase the robustness and reproducibility of the results.

### Definition of regions of interest (ROIs)

ROIs for tractography were defined based on anatomical landmarks on the common template. We defined two ROIs, in the pSTG, and in the pMTG, corresponding to a definition of Wernicke’s area based on anatomical connectivity^9^ and functional relevance for sentence-level processing^10,11^. Taking into account the lack of agreement in the definition of this area^12,13^, we opted for a straightforward anatomical definition of our seed ROIs (described in detail in the Supplementary Methods).

### Probabilistic tractography, generation of connectivity maps in common space

Using diffusion MRI probabilistic tractography, we conducted seed-to-brain tractography from each seed ROI without exclusion or waypoint masks. For each seed ROI in each participant, we obtained a map of the connectivity between each brain voxel and the respective seed region, corresponding to the whole-brain connectivity fingerprint of that seed ROI. In particular, crossing fibre probabilistic tractography^7^ was computed by starting 10,000 streamlines in each voxel of the seed ROI. This algorithm creates a map for the entire brain that represents the number of streamlines that cross each voxel representing the connectivity of this voxel to the seed region. These maps were then logarithmised to obtain a normal distribution of connectivity values, and normalised by the logarithm of the total number of streamlines started in the respective seed ROI, so that we obtained maps with connectivity values ranging between 0 and 1. These maps were spatially registered to the template space with a nearest neighbour interpolation, and smoothed with a Gaussian kernel of 4 mm (FWHM) using the software SPM 12 (University College London, UK). In this way, the voxel values related to the structural connectivity of that particular voxel with the corresponding seed region and will henceforth be called connectivity, and the corresponding maps labelled connectivity maps. Supplementary Figure 1 displays the average connectivity maps across all participants and in each of the language groups.

### Statistical analysis – Machine learning with a Gaussian process classifier

First, we trained a Gaussian process classifier on the complete dataset (N = 134) of the connectivity maps for each of the four seed ROIs (left and right pSTG and pMTG) as implemented in the PRoNTo toolbox, as this was the more adequate procedure for a three-class classification^14^. The cross-validation scheme was implemented by a balanced 10-fold cross-validation from which the average balanced prediction accuracies were obtained. The classifier was run within white matter areas (FA ≥ 0.2) with systematic connections across groups (mean connectivity ≥ 0.2) for each of the seed regions. The double mask was used in all brain analyses to ensure statistics would be computed in white matter regions with connectivity values that are sufficiently sampled by the probabilistic tractography method and therefore might show relevant effect sizes. To prevent the results from being a mere product of group-specific systematic distortions related to macroanatomical differences in brain shape, we computed the mean deformation at each brain voxel from the normalisation field, and excluded in brain areas with strong deformations (Supplementary Figure S3). The performance of the classifier is measured by the average balanced accuracy of the test sets. To obtain the statistical significance of the classification accuracy above chance, a permutation test was run for each of the classifications, with 10,000 permutations^14^. The histograms in Figure 1 (for the left hemispheric ROIs) and Supplementary Figure S2 (for the right hemispheric ROIs) show the probability distribution of the function values of the Gaussian classifier per class, that is language group. Finally, to assess the replicability in independent datasets, we trained the classifier on the connectivity maps pertaining to the larger data set (N = 80) and tested it in the smaller sample (N = 54). To exclude that this effect was due to the use of different scanners, we also ran the classifier in the three subsamples acquired at the same site (German N = 48, Chinese N = 18; taking the unbalanced sample size into account). Again, to verify the statistical significance of the classification accuracy, a permutation test was run for each of the classifications, with 10,000 permutations.

### Statistical analysis – Mass univariate voxel-wise statistics

We then performed spatial voxel-wise statistical comparisons of the normalised connectivity maps of the speakers of each language to identify areas with significant differences in structural connectivity across groups^15^ using SPM 12. This enabled us to identify areas where the connectivity strength was higher in one language group than in the others. Once again, we only considered voxels within the previously defined mask (mean FA ≥ 0.2 and mean connectivity ≥ 0.2). We computed t-tests and report results at p < 0.001 peak-level, p < 0.05 cluster-level family-wise error (FWE) corrected. We both computed direct bidirectional pair-wise comparisons between two languages (Figure 3), and conjunction analysis (Figure 2 and S4). The direct comparisons were performed as a one-sided t-test in the typical implementation of SPM. Besides, we performed a conjunction analysis between the comparisons between each language and the other two. Such an analysis provides us with the particularities of brain structure associated with each language group compared with the other two.

### Tractography from seed ROIs to significant clusters in conjunction analysis

Statistical differences in a specific region obtained in probabilistic tractography do not necessarily reflect local effects. That is why it is fundamental to integrate them in the fibre pathways that connect the seed regions and the significant clusters. In particular, they might result from the overall differences along those fibre pathways. We used probabilistic tractography to establish the course of these pathways crossing both the seed ROI and the region where significant differences had been found. The results of this analysis serve two purposes. On the one hand, the connectivity maps outline the fibre tracts passing by both the posterior temporal cortex and the regions showing stronger connectivity modulation in function of one’s mother tongue. Thus, Supplementary Figure 5 assists us in diagnosing which fibre pathways are involved to a higher degree in a given language.

Furthermore, to assess the magnitude of the connectivity differences across groups in the areas with significant differences, we computed the connectivity strength in a seed to target tractography from the seed ROIs to the respective significant areas. The connectivity values were normalised in the same way as for the creation of the connectivity maps (i.e., logarithmised and divided by the logarithm of the total of streamlines started in the seed region). The plots in the lower row of Figure 2 display the probability distribution of connectivity strength in each of three groups from the seed regions to one of three target areas with significant connectivity differences between groups (left IFG, left ATL, and conjoined significant clusters in the right hemisphere). The white lines in the plots correspond to the percentiles 75, 50, and 25. In these plots, we can see that the magnitude of the differences between the European groups and the Chinese group is comparable with the difference between the two European groups in the significant regions. The Cohens’s d effect sizes of the group differences are shown in a detailed estimation graphic^16^ (see Supplementary Results and Supplementary Figure S9).

### Visualisation of the group averaged tractography

To illustrate the group averaged tractography and relate the areas of significant connectivity differences with the fibre pathways of the language system, we performed an additional group average deterministic tractography as previously used^17,18^ (for details see Supplementary Methods). These representative tractograms were used in Figure 2 and Supplementary Figure 6 to help visualise the fibre tracts where connectivity differences were found.

