## Supplementary Information for "Native language leaves distinctive traces in brain connections"

### Supplementary Methods

**Definition of regions of interest (ROIs).** Concerning the pSTG ROI, we established its anterior border posterior to Heschl's gyrus (transverse temporal gyrus), with the ROI spanning the *planum temporale* to the end of the Sylvian fissure. The pMTG ROI was defined parallel to the pSTG ROI, by transferring its anterior and posterior borders to the MTG (orthogonal to the gyrus). The ROIs did not extend to the fundus of the sulci, only including the superficial half of the gyri to prevent from seeding in deep white matter fibre structures. Analogous to the seed ROIs in the left hemisphere, we also defined two seed ROIs in the right hemisphere taking the contralateral anatomical landmarks as a reference. The seed ROIs were then nonlinearly transformed using the ANTs software from the template space into the native space of single subject. Finally, they were masked with the individual FA map at a threshold of 0.2 to include only anisotropic white matter as seed voxels for tractography.

**Demographic and non-linguistic influence on connectivity differences.** To exclude any possible influence of non-linguistic behavioural differences on the reported results we tested the connectivity measures extracted from the connections with significant differences with a battery of behavioural measures.

**Estimation of the effect size.** To estimate the effect size, we computed Cohen's and the confidence intervals (CI) with bootstrap estimation in R (5000 samples) on the connectivity measures extracted from the connections with significant differences.

**Visualization of the group averaged tractography.** To visualize a group averaged tractography, we normalized the single subject diffusion MRI images to the template image using a deformation map computed from the normalization of the FA images to the template and normalized the intensities across participants. A local diffusion model was then fitted for each voxel using the combination of all normalized diffusion weighted images of 20 participants for each language group distributed over the two samples. Additionally, a model was created including all three languages. We focused on the volume defined by the probabilistic tractography (mean connectivity ≥ 0.2) combined from both seed ROIs and dilated by 2mm. Within this mask, deterministic tractography was started in all voxels (up-sampled voxel size: $1mm^3$) with FA ≥ 0.15. Finally, all streamlines crossing a seed region composed by the combination of the left pSTG and pMTG seed regions were selected. The same procedure was applied to the data from the three language groups and for the data combined over the three languages. Finally, the streamlines connecting the seed regions in the posterior temporal lobe were subdivided in dorsal, ventral and transcallosal connections and coloured differently for both Figure 2 and Figure S6. The streamlines are shown together with a transparent surface of the mean brain template from left lateral and posterior views.

### Supplementary Results

**Differences in connectivity from seed ROIs in the right hemisphere: conjunction analysis.** On the right hemisphere seed ROIs, we only found significant effects in the Chinese group, with stronger connectivity compared to the speakers of the other two languages (Figure S7). Chinese speakers showed a higher connectivity of both the pSTG and pMTG to areas in the contralateral hemisphere, both in the temporal cortex and in subcortical grey matter structures. Additionally, they showed a stronger dorsal connectivity within the right hemisphere.

**Differences in connectivity from seed ROIs in the right hemisphere: pairwise analysis.** We then performed a more detailed pair-wise direct comparison of two languages (Figure S8). First, the comparison between Chinese speakers and English and German speakers respectively showed widespread differences within the right hemisphere and to the contralateral temporal cortex from both seed regions (in the right pSTG and pMTG). Second, when comparing Chinese and English speakers, the former showed stronger connectivity both in the dorsal and ventral fibre pathways of the right hemisphere, besides the cluster in the deep white matter of the left temporal lobe. The English group showed, in turn, no significant clusters with stronger connectivity. When comparing Chinese and German speakers, Chinese speakers exhibited a stronger dorsal connectivity within the right hemisphere in addition to the cluster in the deep white matter of the left temporal lobe. Finally, when comparing English and German speakers, the former yielded higher dorsal connectivity to the right prefrontal cortex from both seed regions. Significant differences were also found in the transcallosal connections. The English group showed, in turn, no significant clusters with stronger connectivity.

**Demographic and on-linguistic influence on connectivity differences.** None of the tested non-lingusitic measures showed a significant correlation with the connectivity measures in the language network. We tested for correlation with age (134), laterality quotient (N=66), foreign language knowledge (N=66), instrument practice (N=66), and phonological memory (N=36). Not all measures were available for all participants and we tested the respective subgroups (N).

**Estimation of the effect size.** The unpaired Cohens'd shows medium to large effect sizes for reported connectivity measures in the language network which support the relevance of the results (Figure S9). The effect sizes and CIs are reported above as: effect size [CI width lower bound; upper bound].

Left anterior temporal ROI (ATL):

Unpaired Cohen's d of German minus English: -0.71 [95CI -1.13; -0.269]

Unpaired Cohen's d of Chinese minus English: -0.613 [95CI -1.04; -0.195]

Left inferior frontal gyrus (IFG):

Unpaired Cohen's d of German minus English: 1.03 [95CI 0.574; 1.43]

Unpaired Cohen's d of Chinese minus English: 0.822 [95CI 0.409; 1.18]

Posterior corpus callosum (PostCC):

Unpaired Cohen's d of English minus Chinese: -0.812 [95CI -1.18; -0.363]

Unpaired Cohen's d of German minus Chinese: -0.54 [95CI -0.888; -0.128]

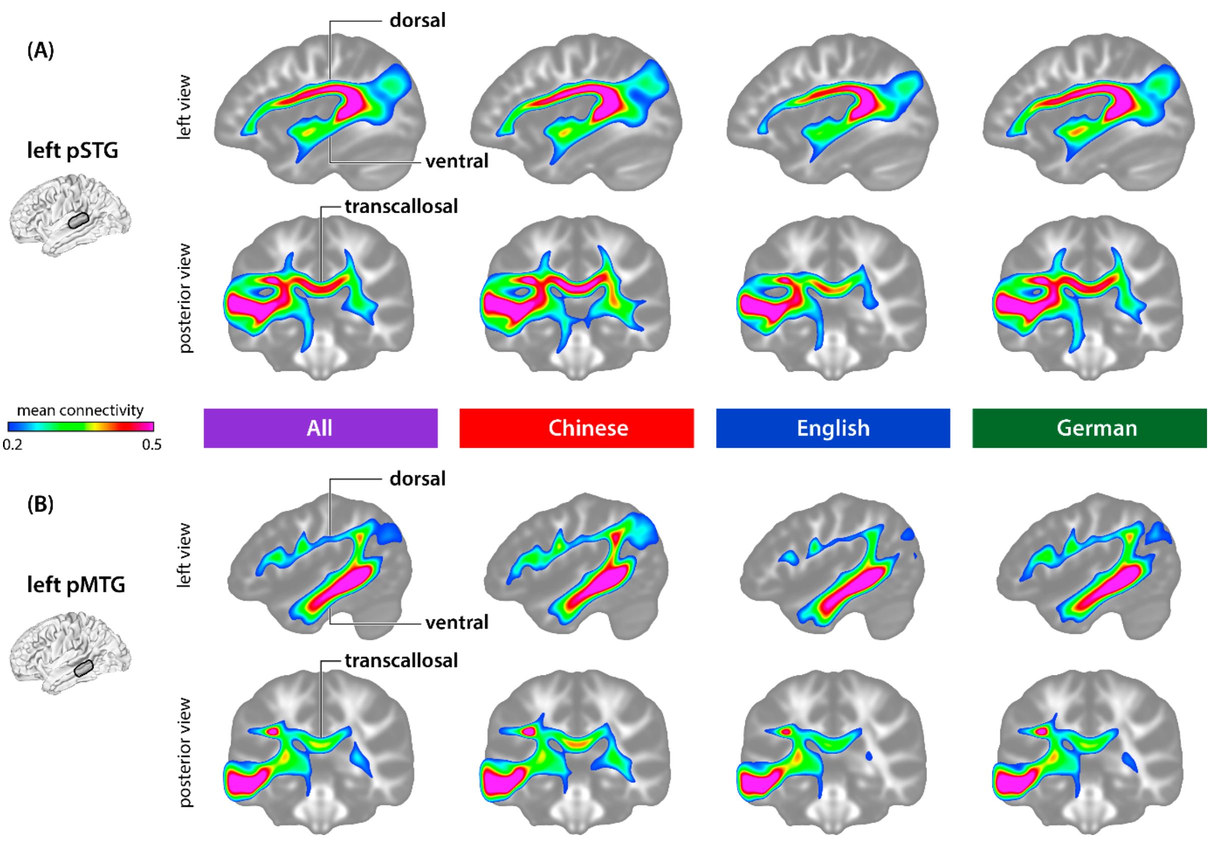


**Figure S1.** Average connectivity maps across all participants and in each of the language groups for (A) seed in left pSTG and (B) seed in pMTG seed regions).

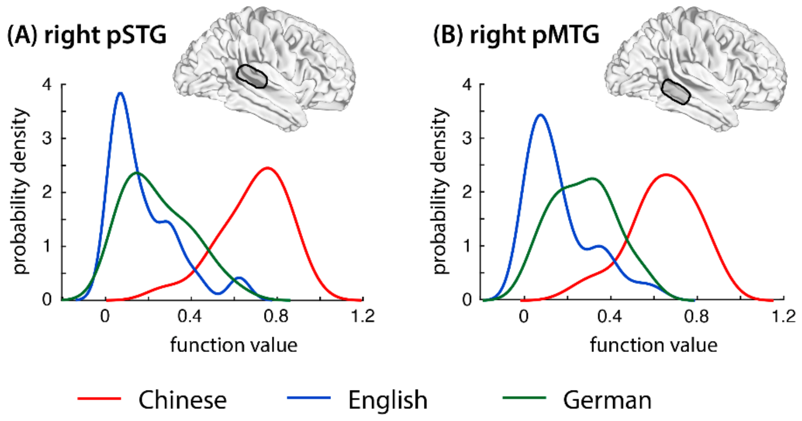


**Figure S2.** Performance of the classifier. Performance of the classifier on the connectivity map of the two seed ROIs in the right hemisphere, (A) right posterior superior temporal gyrus (pSTG) and (B) right middle temporal gyrus (pMTG). Performance for the three different languages is colour coded red for Chinese, blue for English and green for German.

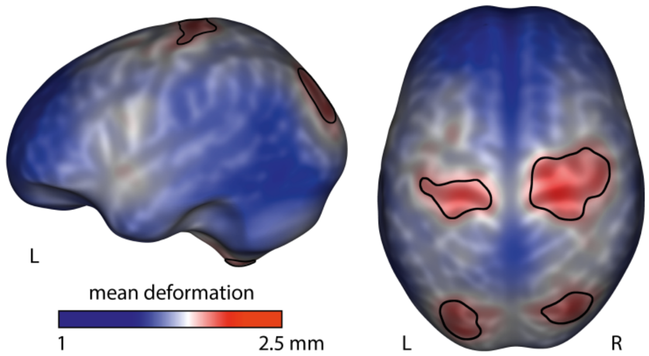


**Figure S3.** Average magnitude of the deformation field of all participants. Excluded regions with strong deformations above 75% of the maximum deformation are marked.

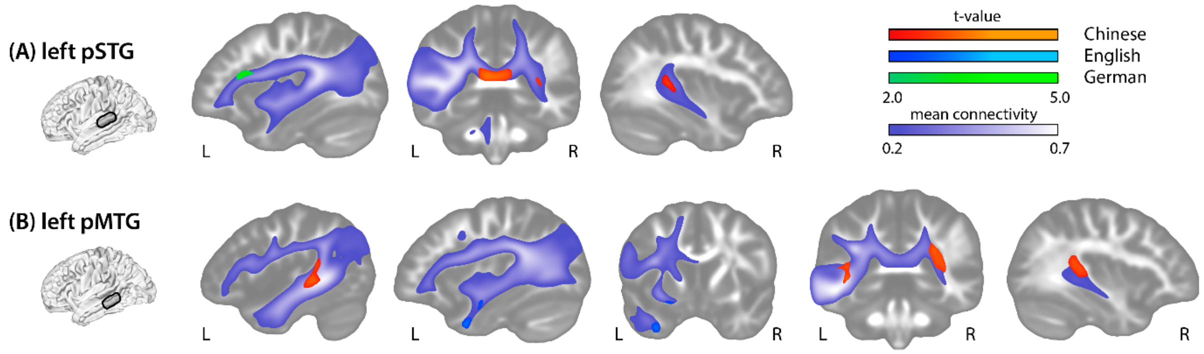


**Figure S4.** Cross-linguistic differences in connectivity strength from (A) left pSTG and (B) pMTG in the conjunction analysis. The white matter mask (FA>0.2) with a mean connectivity > 0.2 used for statistical analysis is shown with a purple scale.

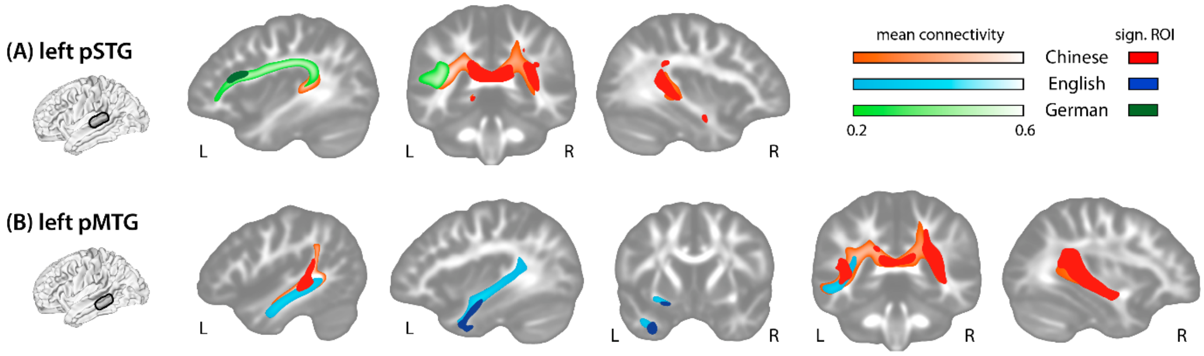


**Figure S5.** Identification of fibre tracts with significant differences in connectivity with probabilistic tractography from seed regions in the (A) left pSTG and (B) left pMTG. Languages are colour coded: red for Chinese, blue for English and green for German.

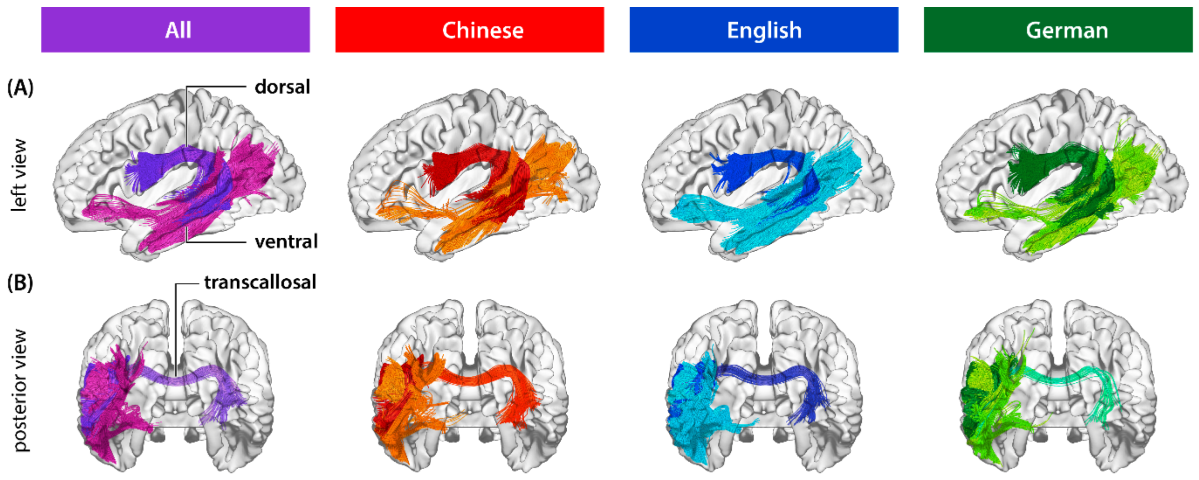


**Figure S6.** Representative tractograms from left pSTG and pMTG seed regions across all participants and in each of the language groups. (A) left view, displaying the dorsal and ventral fibre tracts. (B) posterior view visualizing the transcallosal connection.

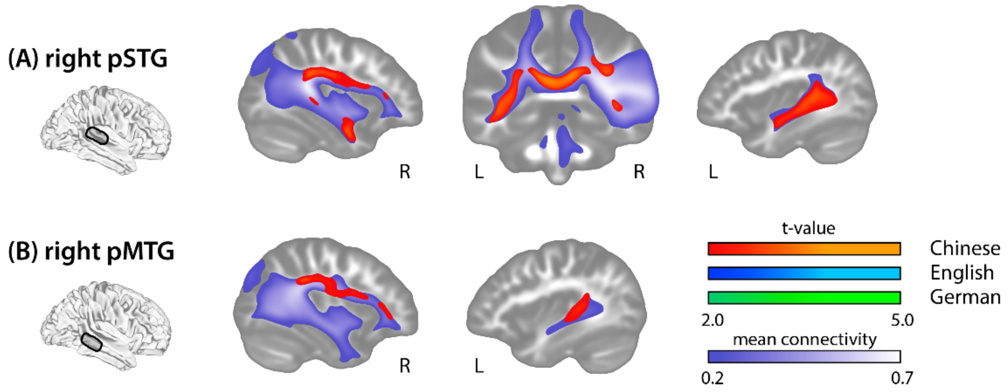


**Figure S7.** Cross-linguistic differences in connectivity strength from (A) right pSTG and right (B) pMTG in the conjunction analysis.

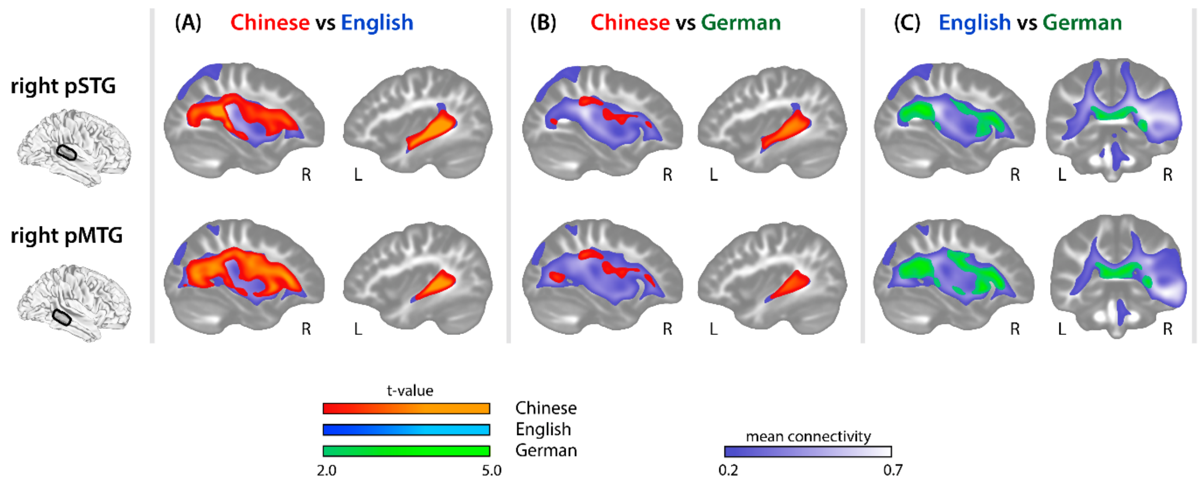


**Figure S8.** Pair-wise connectivity differences between each pair of the three languages from right seed regions: (pSTG and pMTG), (A) Chinese vs English, (B) Chinese vs German, (C) English vs German.

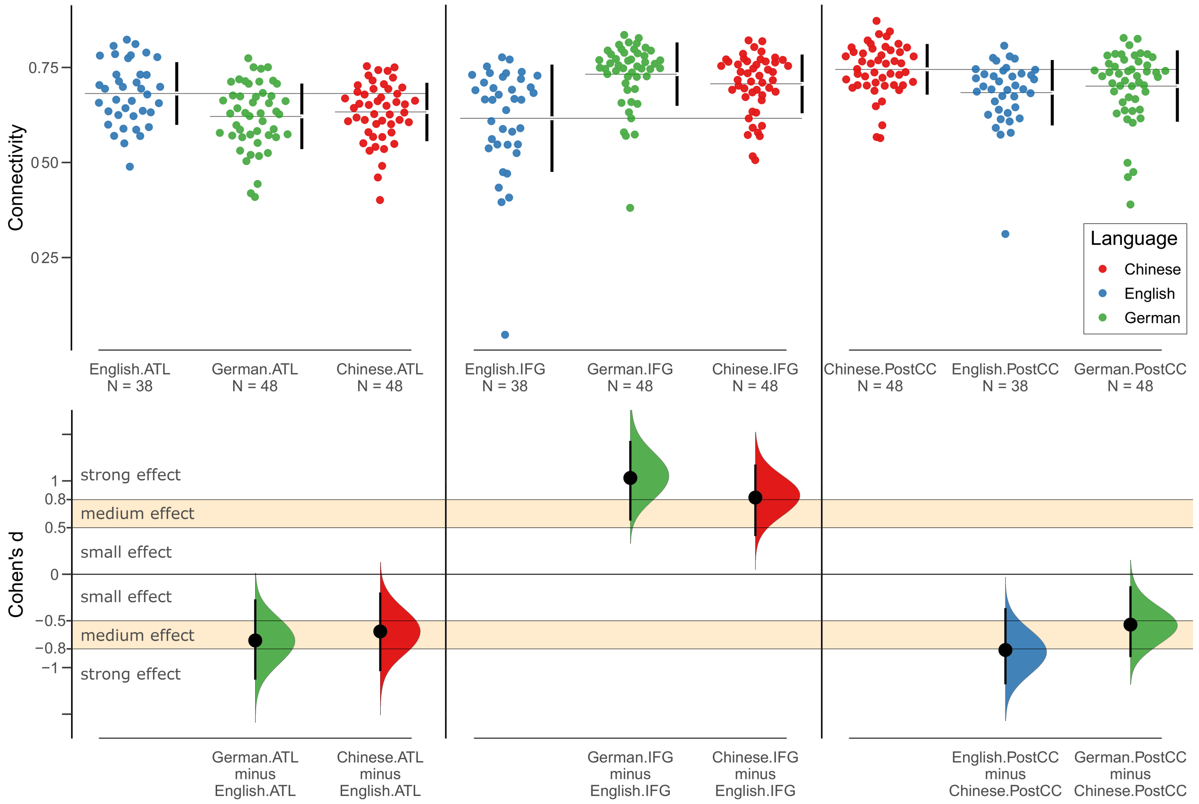


**Figure S9.** Effect size estimation. Top row: estimation graphics of the probability distribution of connectivity strength in each of three groups from the seed regions to one of three target areas with significant connectivity differences between groups (left IFG, left ATL, and conjoined significant clusters in the right hemisphere) as shown Figure 2. Bottom row: The Cohen's d between the groups are shown in the Gardner-Altman estimation plots. The mean difference is plotted on a floating axes as a bootstrap sampling distribution. The mean difference is depicted as a dot; the 95% confidence interval is indicated by the ends of the vertical error bar.
